# Causal role for sleep-dependent reactivation of learning-activated sensory ensembles for fear memory consolidation

**DOI:** 10.1101/2020.04.30.070466

**Authors:** Brittany C. Clawson, Emily J. Pickup, Amy Enseng, Laura Geneseo, James Shaver, John Gonzalez-Amoretti, Meiling Zhao, A. Kane York, Sha Jiang, Sara J. Aton

**Author notes:** Corresponding author: Dr. Sara J. Aton, University of Michigan, Department of Molecular, Cellular, and Developmental Biology, 4268 Biological Sciences Building, 1105 N. University Ave, Ann Arbor, MI 48109.

## Abstract

Learning-activated engram neurons play a critical role in memory recall. An untested hypothesis is that these same neurons play an instructive role in offline memory consolidation. Here we show that a visually-cued fear memory is consolidated during post-conditioning sleep in mice. We then use TRAP (targeted recombination in active populations) to genetically label or optogenetically manipulate primary visual cortex (V1) neurons responsive to the visual cue. Following fear conditioning, mice respond to activation of this visual engram population in a manner similar to visual presentation of fear cues. Cue-responsive neurons are selectively reactivated in V1 during post-conditioning sleep. Mimicking visual engram reactivation optogenetically leads to increased representation of the visual cue in V1. Optogenetic inhibition of the engram population during post-conditioning sleep disrupts consolidation of fear memory. We conclude that selective sleep-associated reactivation of learning-activated sensory populations serves as a necessary instructive mechanism for memory consolidation.

## Introduction

Experiences during wake influence neural activity patterns during sleep. For example, hippocampal place cells activated during environmental exploration in wake show higher firing rates (reactivation)^1^ and/or similar sequences of activity (replay)^2–6^ during subsequent sleep. This phenomenon has been observed in multiple brain regions, multiple species, and following a wide range of experiences^7–13^. Since sleep loss has a disruptive effect on many forms of memory^14^, replay and reactivation may play an instructive role in sleep-dependent memory consolidation^14,15^. To test this, prior work has disrupted network-wide activity during specific sleep oscillations^16–19^ or disruption of activity in genetically-defined cell types across specific phases of sleep^20–23^ - but not the specific neurons activated during learning itself. Recent work has emphasized the essential role of engram neurons in memory recall ^24,25^. To date however, no studies have applied this technology to the question of sleep-dependent memory consolidation.

Here we test the necessity of sleep-specific engram neuron reactivation for memory consolidation. We describe a form of visually-cued fear memory in mice, which is encoded by single trial conditioning (pairing presentation of an oriented grating visual stimulus with an aversive foot shock) and dependent on post-conditioning sleep. Post-conditioning, the mice behaviorally discrimination between conditioned and neutral visual cues, leading to a selective fear memory. This discrimination is disrupted by post-conditioning sleep deprivation. Using this paradigm, we take advantage of recently developed genetic tools to selectively manipulate orientation-selective (i.e., cue-activated) primary visual cortex (V1) neurons. We find that these cue-activated visual engram neurons are selectively reactivated during sleep in the hours following visually-cued fear conditioning. Optogenetic stimulation of these neurons in awake behaving mice generates a percept of the fear cue, which is sufficient to drive both fear learning and recall. A period of rhythmic optogenetic activation of cue-activated neurons is sufficient to drive functional plasticity - increasing representation of the cue orientation in surrounding V1 neurons - and their optogenetic inhibition reduces cue orientation preference. Finally we show that selective sleep-targeted inhibition of cue-activated V1 neurons during post-conditioning sleep is sufficient to disrupt consolidation of visually-cued fear memory. Based on these findings, we conclude that neurons that are selectively activated in sensory cortical areas during learning play an instructive role in subsequent sleep-dependent memory consolidation.

## Results

### Visually-cued fear memory consolidation is disrupted by sleep deprivation

We first tested the role of sleep in consolidating fear memory associated with a specific visual cue. At ZT0, wild-type mice underwent visually-cued fear conditioning in a novel arena (context A), in which three 30-s presentations of phase-reversing gratings (of a specific orientation X°, shown on 4 LED monitors surrounding the arena) coterminated with a 2-s foot shock. Mice were then returned to their home cage and either allowed *ad lib* sleep for the next 12 h, or sleep deprived (SD) for 6 h followed by 6 h of *ad lib* recovery sleep. At ZT12, fear memory for the visual shock cue was assessed in a distinct novel context B. During two separate tests, mice were exposed to gratings of either the same orientation (*i.e*., shock cue X°) or a different orientation (neutral cue Y°) (**Figure 1a**). As shown in **Figure 1b**, mice allowed *ad lib* sleep showed significantly higher freezing responses during presentation of the shock cue than presentation of the neutral cue (two-way RM ANOVA, effect of sleep condition: *p* = 0.014, effect of cue orientation: *p* < 0.001, sleep condition × orientation interaction: *p* = 0.047). Both sleeping and SD mice discriminated between the shock and neutral cue (*p* < 0.0001 for sleep, *p* = 0.03 for SD, Holm-Sidak *post hoc* test), however, SD mice displayed significantly less freezing to the shock cue than mice allowed *ad lib* sleep (*p* = 0.002, Holm-Sidak *post hoc* test). To compare discrimination between cues, a discrimination index was calculated. Both freely sleeping and SD mice showed discrimination that differed from chance values, however this effect was far clearer in mice allowed *ad lib* sleep (Wilcoxon signed rank test; Sleep: *p* = 0.0003, SD: *p* = 0.049). **Figure 1** shows data for both female and male mice (males - filled symbols, females - open symbols; a breakdown by sex is provided in **Extended Data Figure S1**). Both sexes displayed discrimination between shock and neutral cues when allowed *ad lib* sleep (*p* = 0.001 and *p* = 0.007 for males and females respectively, Holm-Sidak *post hoc* test) and impairment when sleep deprived (*N.S*. for shock vs. neutral, Holm-Sidak *post hoc* test). Both sexes showed significant discrimination from random chance only when sleep was allowed (*p* = 0.016 for both male and female freely-sleeping mice, *N.S*. for male and female SD mice, Wilcoxon signed rank test). Thus, for subsequent analysis, both sexes were used.

**Figure 1.**
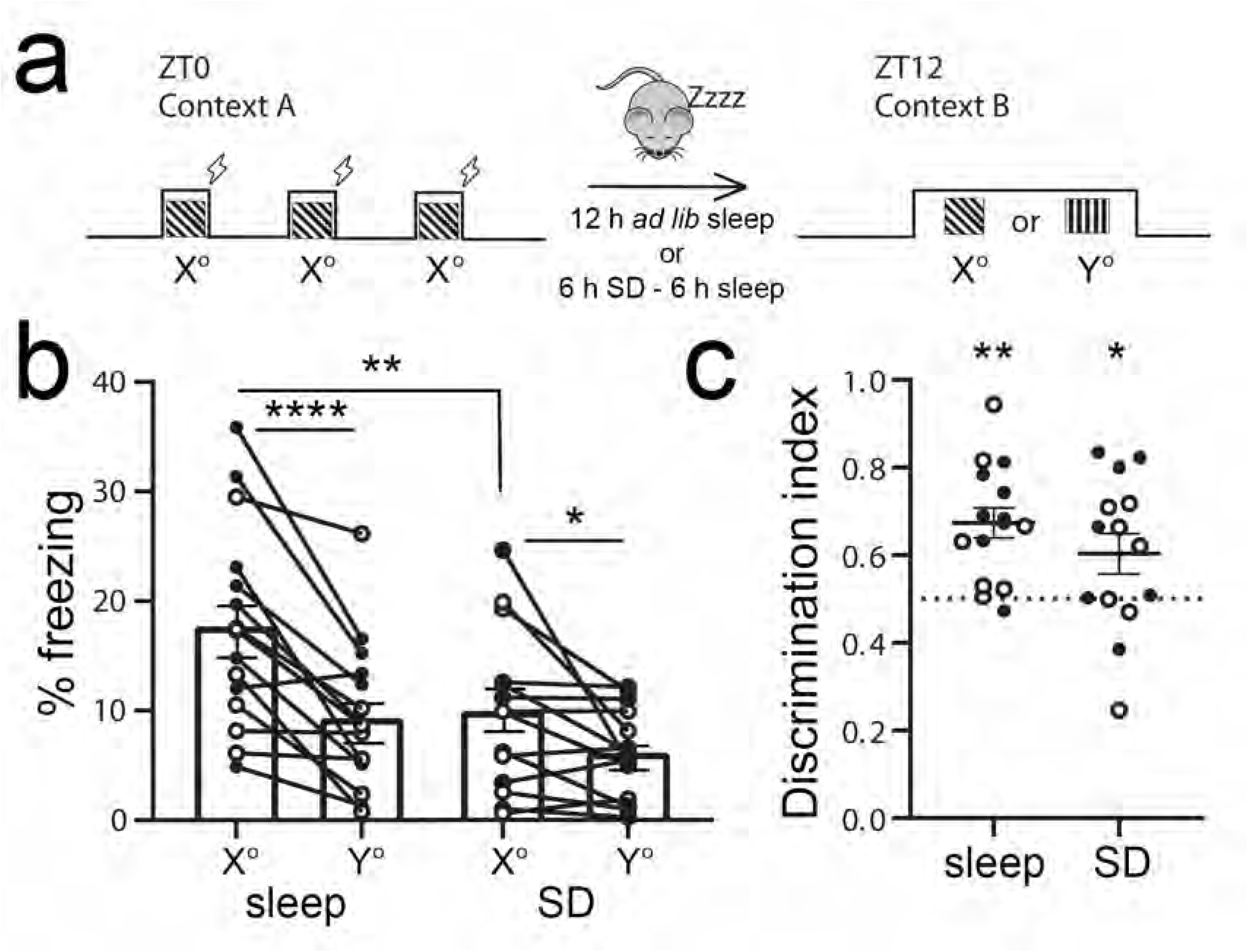
Consolidation of visually-cued fear memory is enhanced by post-conditioning sleep. **(a)** At ZT0, mice underwent three stimulus-shock pairings in context A. After either 12 h of *ad lib* sleep or 6 h sleep deprivation (SD) followed by 6 h *ad lib* sleep, mice were exposed to the shock cue (X° grating) and a neutral cue (Y° grating) in context B. **(b)** Freezing behavior of the mice during the ZT12 test (Sleep: *n* = 15 - sleep, SD: *n* =14; males - solid symbols, females - open symbols). Mice allowed to sleep froze significantly more to the shock cue than mice who were sleep deprived (** indicates *p* = 0.002, Holm-Sidak *post hoc* test). Both freely-sleeping and SD mice showed higher freezing in response to the shock cue (**** indicates *p* < 0.0001, * indicates *p* = 0.03, Holm-Sidak *post hoc* test; two-way RM ANOVA: main effect of sleep condition, *F* = 6.9, *p* = 0.014, main effect of orientation, *F* = 28.5, *p* < 0.001, sleep x orientation interaction, *F* = 4.4, *p* = 0.047). **(c)** Freezing behavior quantified a discrimination index [X°/(X°+Y)] for each mouse and compared to chance performance (* indicates *p* = 0.049, ** indicates *p* = 0.0003, Wilcoxon signed rank test vs. chance).

The neural circuitry underlying visually-cued fear memory could be altered by sleep. Prior work from our lab has shown that presentation of oriented gratings can initiate response plasticity in neurons of the lateral geniculate nucleus (LGN) and V1, which are consolidated during subsequent sleep^20,26–28^. However, prior studies of auditory-cued fear have shown conflicting results on the necessity of post-conditioning sleep for consolidation^29–32^. We hypothesized that these discrepancies could be due to differences in timing of either training or testing (or both) between studies. To test this, we performed a time course of fear memory testing for mice conditioned at ZT0. We found that freely-sleeping mice showed differential visual cue discrimination when tested 12, 24, and 36 h after visually-cued fear conditioning - with clear discrimination between shock and neutral cues seen at ZT12 time points (12 and 36 h post-conditioning, *p* = 0.001 and *p* < 0.001 respectively, Holm-Sidak *post hoc* test; **Extended Data Figure S2**) but not at ZT0 (24 h post-conditioning; *N.S*., Holm-Sidak *post hoc* test). At no time point did SD mice discriminate between shock and neutral cues (all *N.S*., Holm-Sidak *post hoc* test). Together these data suggest that visually-cued fear memory consolidation is sleep-dependent, while fear recall may show diurnal rhythmicity.

### Targeted recombination in activated populations (TRAP) targets orientation-selective, fear cue-activated V1 neurons

To characterize and manipulate activity in V1 neuronal populations activated by oriented grating cues (*i.e*., putative visual engram neurons), we used previously described techniques for TRAP^33^. cfos-CRE^ER^ mice were crossed to mice expressing tdTomato in a cre-dependent manner (*cfos::tdTom*). The mice were presented with either an oriented grating (X°) or a dark screen stimulus for a 30-min period (**Figure 2a**). Immediately following this presentation, mice were administered tamoxifen and housed in complete darkness for the next 3 days (to prevent additional visually-driven recombination in V1). V1 tdTomato expression (quantified 11 days following tamoxifen administration) was significantly higher in mice exposed to gratings; dark screen presentation induced very low levels of V1 expression (**Figure 2b-c**; nested t-test, *p* = 0.0001).

**Figure 2.**
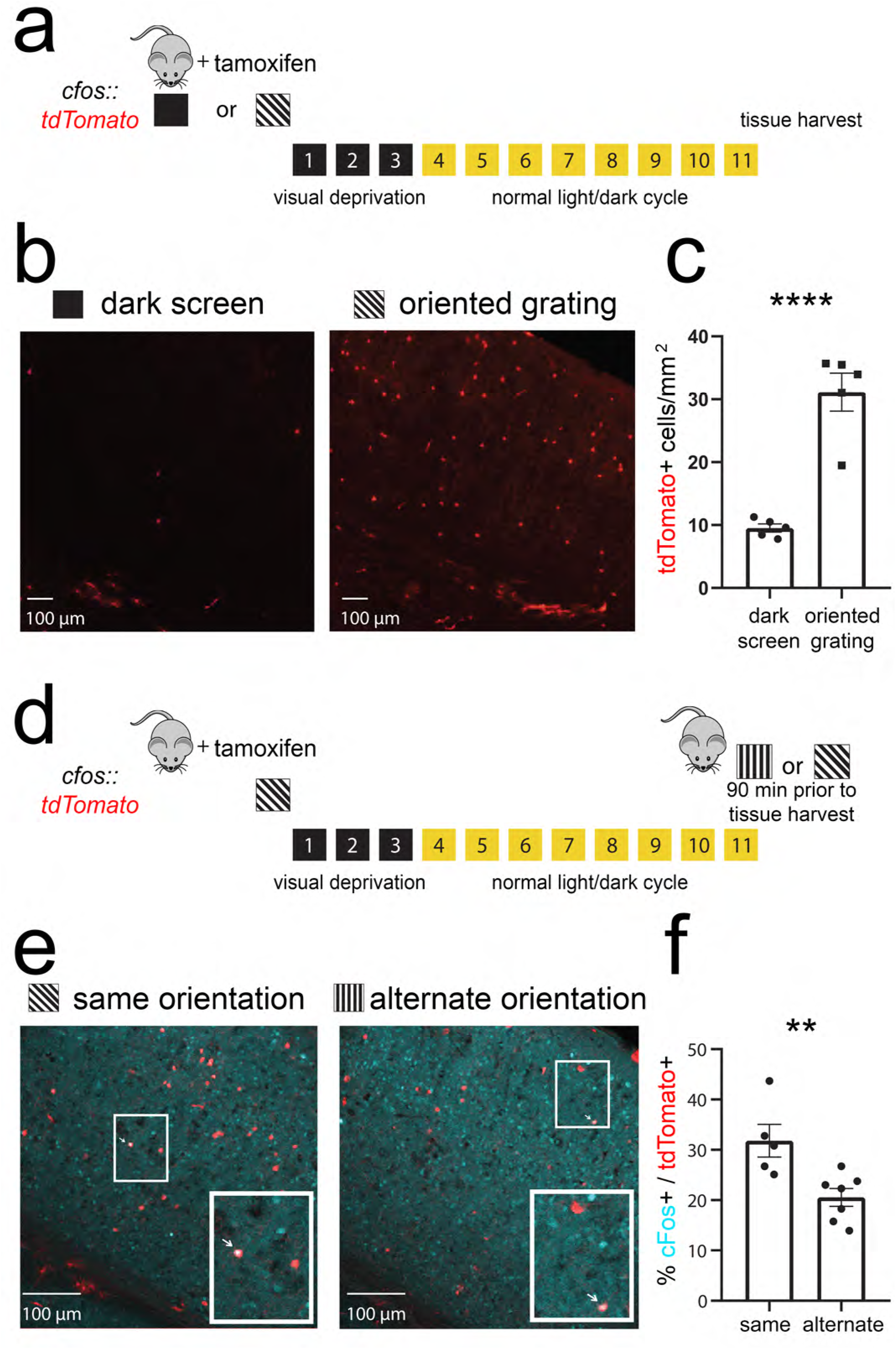
TRAP labels orientation-selective V1 ensembles. **(a)** *cfos::tdTom* mice were presented with either a dark screen or an oriented grating (X°) and were then injected with tamoxifen prior to 3 d of housing in complete darkness. **(b-c)** Representative V1 tdTomato labelling quantified 11 d after tamoxifen administration. **** indicates *p* = 0.0001 (*t* = 7.07, *DF* = 8) for dark screen vs. X°, nested t-test (*n* = 5 mice/condition) **(d)** Prior to tissue harvest, mice were either re-exposed to gratings of the same orientation (X°) or an alternate orientation (Y°). **(e-f)** Representative images showing overlap of tdTomato (red) and cFos protein (cyan). An example of colocalization within a neuron (quantified in **f**) is indicated with a white arrow for each image in the inset. ** indicates *p* = 0.009 (*t* = 3.22, *DF* = 10), nested t-test (*n* = 5 mice for X°, *n* = 6 mice for Y°).

To test the orientation selectivity of X°-activated TRAPed neurons, mice were presented with either the same oriented grating (X°) or an alternate oriented grating (Y°) prior to sacrifice (**Figure 2d**). TRAPed V1 neurons show a significantly higher percent cFos expression following re-exposure to the same orientation than following exposure to a different orientation (X°- 32 ± 3% vs. Y°- 21 ± 2%; *p* = 0.009, nested t-test; **Figure 2e-f**). This level and specificity of cFos overlap is comparable to that reported for auditory stimuli in cochlear nuclei (Guenthner et al 2013). Together these data suggest that TRAP provides genetic access to orientation-selective V1 neurons activated by oriented grating stimuli.

### Optogenetic activation of TRAPed V1 neurons generates a orientation-specific percept

To further test the cue selectivity of recombination in neurons activated by an oriented grating (X°), and to test the behavioral significance of activity in this neuronal population, we expressed ChR2 in X°-activated TRAPed neurons (*cfos::ChR2*). As shown in **Figure 3a**, *cfos::ChR2* mice implanted with bilateral V1 optic fibers were presented with a single oriented grating (X°) for TRAP as described above; 11 days later, one of two variants of the visually-cued fear conditioning were performed. The first subset of mice were conditioned at ZT0 using rhythmic (1Hz) optogenetic activation of TRAPed V1 neurons (rather than oriented grating presentation) as a cue for foot shock (**Figure 3b**). Mice were returned to their home cages, allowed *ad lib* sleep, and tested at ZT12 in a dissimilar context. At this point, mice were presented with oriented gratings of the same orientation used for TRAP (X°) and a different (Y°) orientation. Presentation of X° elicited significantly greater freezing responses than presentation of Y° (ratio paired t-test, *p* = 0.008; Wilcoxon signed rank test vs. chance, *p* = 0.02) (**Figure 3c**).

**Figure 3.**
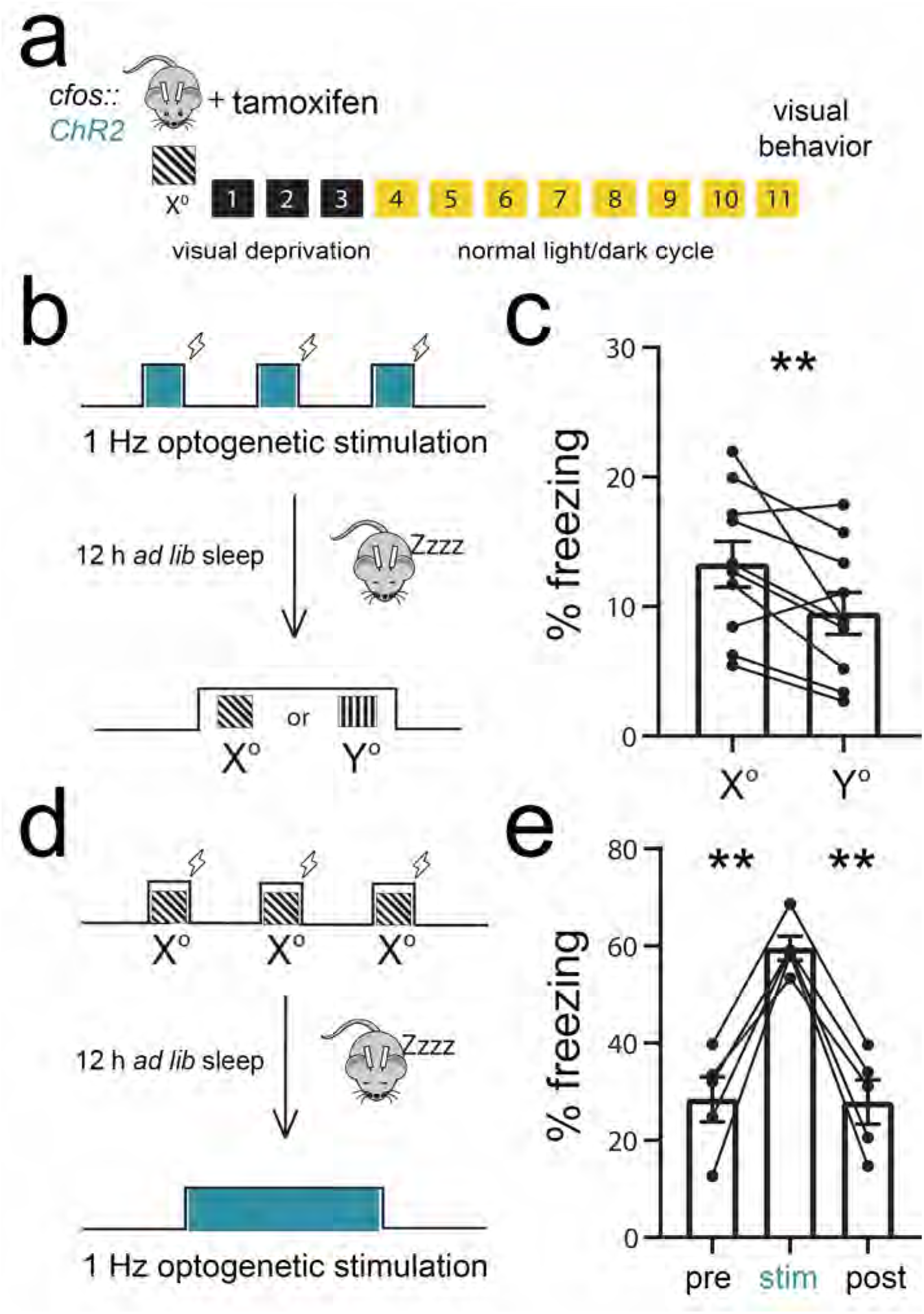
Optogenetic stimulation of TRAPed V1 neurons mimics visual experience. **(a)** *cfos::ChR2* mice with bilateral V1 fiber optics had recombination induced to a specific angle (X°). 11 days later, visual behavior was run. **(b-c)** At ZT 0, the mice received bilateral V1 optogenetic stimulation paired with foot shocks in lieu of the oriented grating visual stimuli used for cued conditioning in **Figure 1**. At ZT 12, the mice were presented with the same oriented grating used for TRAP (X°) and an alternate orientation (Y°). Optogenetically-cued conditioning resulted in higher subsequent cued freezing responses to X° relative to Y° (*n* = 10 mice; *p* = 0.008 [*t* = 3.38, *DF* = 9], ratio paired t-test). **(d-e)** At ZT0, a second cohort of mice underwent visually-cued fear conditioning to the same orientation as the TRAPed ensemble. At ZT12, the mice received optogenetic stimulation in place of a visual test. Freezing behavior was higher during optogenetic stimulation than before or after stimulation (*n* = 5 mice; pre vs. stim - *p* = 0.003 [*t* = 7.30, *DF* = 4, stim vs. post - *p*=0.003 [*t* = 7.85, *DF* = 4], Holm-Sidak *post hoc* test, one-way RM ANOVA).

In a second set of experiments (**Figure 3d**), mice underwent visually-cued fear conditioning at ZT0, using X° gratings as the shock cue. At ZT12, they were placed in a dissimilar context, where after a delay they received bilateral 1 Hz optogenetic stimulation of TRAPed V1 neurons. These mice showed significantly greater freezing behavior during optogenetic stimulation than before and after stimulation (**Figure 3e**; *p* = 0.003 for each Holm-Sidak *post hoc* test). Both of these results indicate that optogenetic activation of the X°-activated TRAPed V1 ensemble is sufficient to generate a percept of the X° visual cue, consistent with recent data^34^. Moreover, these data demonstrate that optogenetically-activated V1 neurons can substitute behaviorally as a visual cue for either encoding or recalling fear memory. Together, this suggests that activity of the X°-activated TRAPed ensemble in V1 constitutes an engram for the visual cue.

### Orientation-selective V1 ensembles are reactivated during post cued fear conditioning sleep

Since sleep facilitates consolidation of visually-cued fear memory, and the TRAPed ensemble provides cue-selective information, we next evaluated whether the TRAPed population is selectively activated during post-conditioning sleep. We again expressed tdTomato in TRAPed X°-activated neurons (*cfos::tdTom*). As shown in **Figure 4a**, these mice were presented with X° to induce tdTomato expression, and 11 d later were cue conditioned using either the same X° oriented grating stimulus, or a dissimilar Y° stimulus. They were then returned to their home cage and allowed *ad lib* sleep over the next 4.5 h, at which point they were sacrificed for V1 cFos immunostaining. When X° was used as the fear conditioning cue, 34 ± 2% of tdTomato-expressing V1 neurons showed expression of cFos after subsequent sleep (**Figure 4b**) - a level similar to that seen after same-orientation grating exposure (**Figure 2e**). When mice were instead conditioned using Y° as the shock cue, the percent overlap was significantly lower (26 ± 1%). These data suggest V1 neurons activated by a visually-cued learning experience are more likely to remain active during post-learning sleep, consistent with observations of ensemble reactivation in V1 following other types of learning^7^. Thus sleep-associated V1 ensemble reactivation could serve as a plausible substrate underlying visually-cued fear memory consolidation.

**Figure 4.**
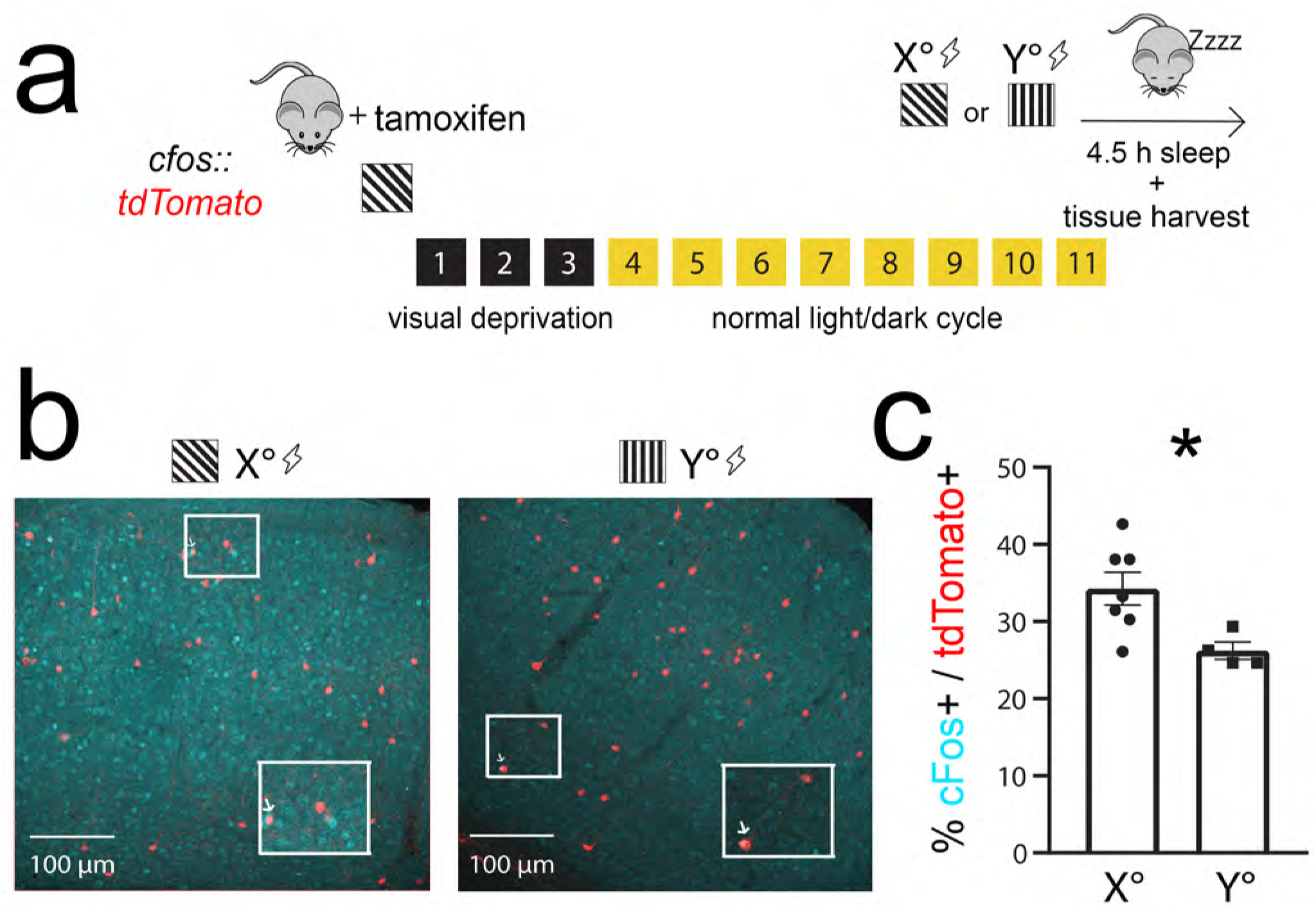
TRAPed V1 neurons selective for the conditioned stimulus are reactivated in post-conditioning sleep. **(a)** *cfos::tdTom* mice had recombination induced to a specific angle (X°). 11 days later, they were cue conditioned to either the same angle as induction (X°; *n* = 7 mice) or an alternate angle (Y°; *n* = 4 mice). All mice were allowed 4.5 h of post-conditioning *ad lib* sleep prior to tissue harvest. **(b-c)** Representative images showing overlap of cFos expression (cyan) with tdTomato (red). The boxed region is magnified as an inset with an arrow indicating an overlapping neuron. Expression of cFos in tdTomato-labelled cells was greater for mice conditioned to the same orientation used for TRAP labelling (* indicates *p* = 0.025 [*t* = 2.69, *DF* = 9], nested t-test).

### Rhythmic offline reactivation of orientation-selective V1 ensembles induces plasticity and alters representation of orientation in V1

To test whether sleep-associated reactivation of orientation-selective neurons could impact the representation of orientation across V1, we tested how rhythmic optogenetic activation of X°-activated TRAPed neurons affected surrounding V1 neurons’ response properties. We recorded neuronal firing patterns and visual responses in V1 from anesthetized *cfos::ChR2* mice before, during and after a period of rhythmic (1 Hz) light delivery. We first generated tuning curves to assess orientation preference for each V1 neuron, measuring firing rate responses to a series of 8 different oriented gratings. This orientation preference test was followed by a 20-30 min period without optogenetic stimulation, a second orientation preference test, a 20-30 min period of 1 Hz optogenetic stimulation, and then a final orientation preference assessment (**Figure 5a**).

**Figure 5.**
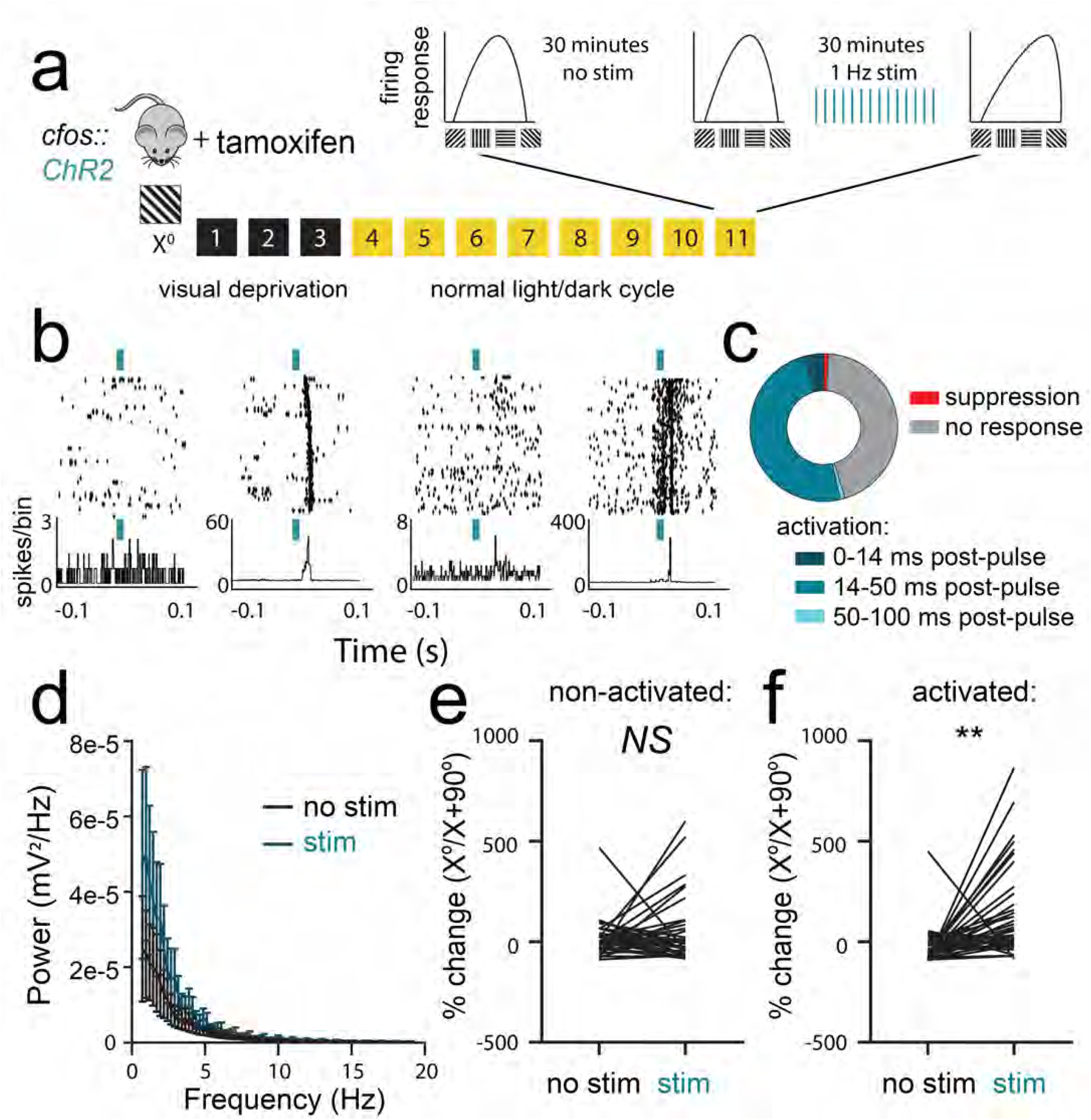
Offline reactivation of orientation-selective TRAPed V1 neurons alters orientation representations in V1. **(a)** *cfos::ChR2 mice* were presented with an oriented grating (X°) for TRAP. 11 days later, orientation tuning was measured repeatedly for V1 neurons recorded from anesthetized mice: at baseline, after a 20-30 min period without optogenetic stimulation, and after a 20-30 min period with 1 Hz light delivery. **(b)** Representative rasters and perievent histograms for 4 simultaneously-recorded neurons, showing diverse firing responses during optogenetic stimulation. **(c)** The majority of stably-recorded V1 neurons were reliably activated following light pulses, with variable lag times. A small proportion were inhibited by light delivery, and the remaining neurons were not affected (*n* = 62 neurons from 5 mice, total). **(d)** Power spectra for V1 LFPs showed no significant effect on ongoing rhythmic activity (*N.S*., K-S test, *n* = 5 mice) **(e-f)** After optogenetic stimulation, neurons that were not activated following light pulses showed no change in orientation preference (*N.S*., nested t-test, *n* = 32 neurons from 5 mice). In contrast, activated neurons showed increased firing rate responses for gratings of the same orientation (X°) used for TRAP. (** indicates *p* = 0.002 [*t* = 3.27, *DF* = 62], nested t-test, *n* = 30 neurons from 5 mice).

V1 neurons showed heterogeneous firing responses during rhythmic optogenetic stimulation (**Figure 5b**). A small fraction of the recorded neurons (4%) were activated immediately following initiation of the 10-ms light pulses, 1% were significantly inhibited, and 1% showed only long-latency (more than 200 ms) excitatory responses. The remaining recorded neurons were either unaffected by optogenetic stimulation (44%) or showed consistent activation 14-50 ms after light pulses (49%), suggesting these neurons receive excitatory input from the optogenetically-stimulated population (**Figure 5c**). Rhythmic activation of the X°-activated V1 population did not significantly alter the V1 local field potential (LFP) power spectrum (**Figure 5d**, *N.S*., K-S test).

To assess how optogenetic reactivation of the X°-activated TRAPed population affects response properties in surrounding V1 neurons, orientation tuning curves for well-isolated and stably-recorded neurons were compared before vs. after optogenetic stimulation. While orientation preference for X° (vs. X+90°) was stable across 20-30 min period without optogenetic stimulation, a similar period of 1 Hz light delivery caused a selective shift in orientation preference across V1 toward the orientation of the TRAPed population. Shifts in orientation preference towards the orientation of the TRAPed ensemble (X°) were greater for those neurons that showed consistent excitatory responses 20-50 ms following light pulses, relative to neurons that did not show these responses (*N.S*. for non-activated neurons, vs. *p* = 0.002 for activated neurons, nested t-test; **Figure 5e,f**). Critically, this shift is similar to that seen in V1 after presentation of oriented gratings, followed by a subsequent period of *ad lib* sleep^20,27,28^.

### Sleep-associated reactivation of orientation-selective V1 neurons is necessary for consolidation of visually-cued fear memory

Because reactivation of orientation-selective V1 populations occurs during post-visually-cued conditioning sleep, and is sufficient to induce changes in orientation representations in V1, we next tested the necessity of sleep-associated ensemble reactivation for consolidation of visually-cued fear memory. To assess how inhibition of the X°-activated TRAPed population affects firing in surrounding V1 neurons, we expressed ArchT in cfos-CRE^ER^ mice (*cfos::Arch*). We recorded spontaneous activity and visual responses in V1 neurons in anesthetized mice before and during a period of optogenetic inhibition (**Figure 6a**). Periodic inhibition (cycles of 5 s light delivery, followed by a 0.5 s ramp off, and 1 s off) led to heterogeneous changes in spontaneous firing (**Figure 6b-c**), with 34% showing no response (± 0-5% change in firing rate), 21% activated (> 5% increase in firing rate) and 34%, 9%, and 2%, respectively, inhibited slightly (6-33% decrease in firing rate), moderately (34-66% decrease), or strongly (67-100% decrease). Inhibition did not affect V1 LFP power spectra (**Figure 6d**, *N.S*., K-S test). Inhibition during presentation of oriented gratings led to a significant decrease in orientation preference for X° in inhibited neurons (**Figure 6f**, *p* = 0.007, nested t-test). There was no significant shift in non-inhibited neurons (**Figure 6e**, *N.S*., nested t-test). Together, these data indicate that inhibition of the TRAPed ensemble leads to changes in orientation representation across the population, without grossly disrupting network activity across V1.

**Figure 6.**
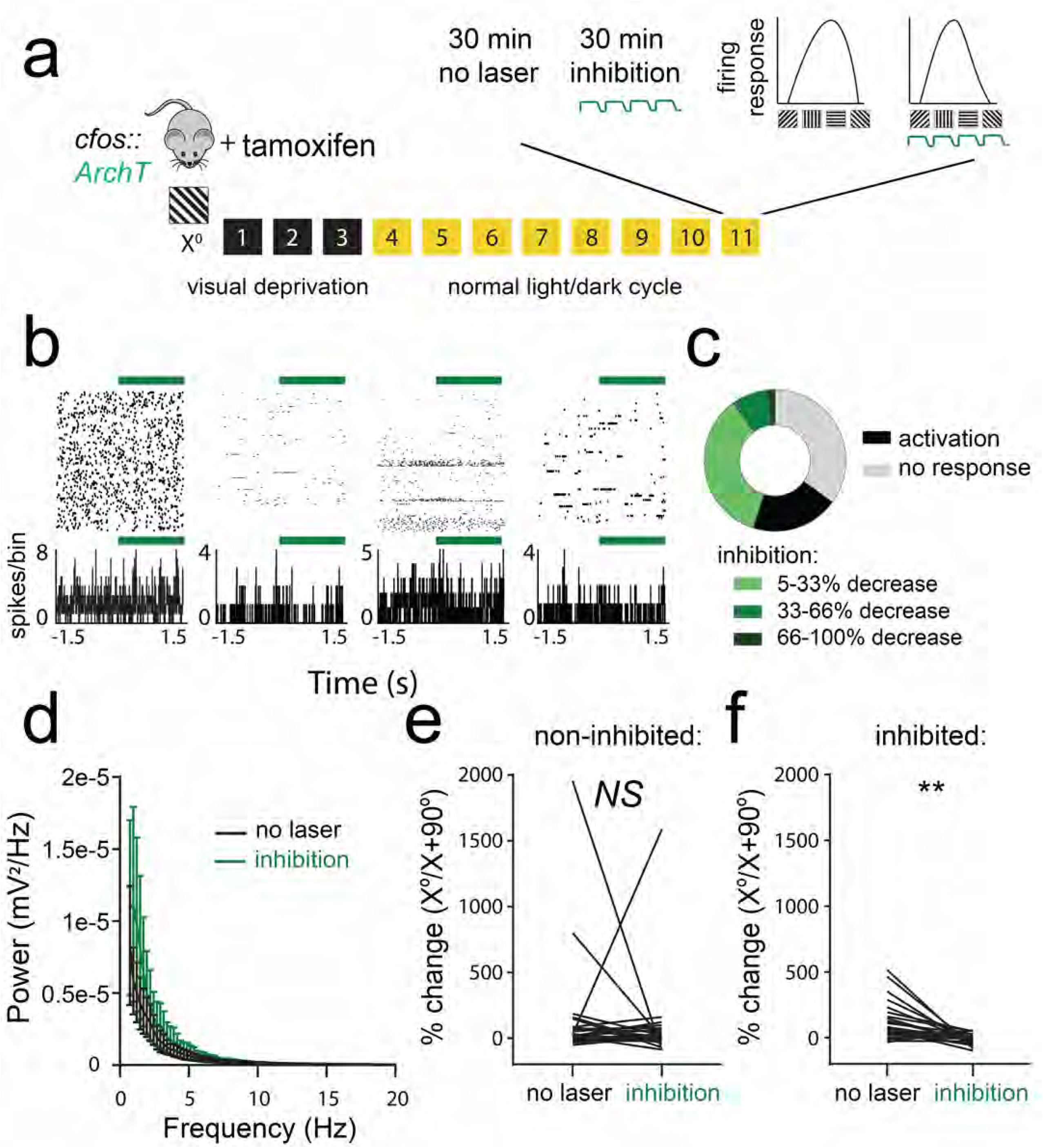
Optogenetic inhibition of orientation-selective TRAPed V1 ensembles alters orientation preference in surrounding V1 neurons. **(a)** *cfos::ArchT* mice were presented with an oriented grating (X°) for TRAP. 11 days later, V1 neurons were recorded from anesthetized mice across 30 minutes of optogenetic inhibition, and 30 minutes without inhibition. Afterward, orientation preference was assessed without optogenetic inhibition (no laser) and with inhibition. **(b)** Representative rasters and perievent histograms for 4 simultaneously-recorded neurons, showing diverse firing responses during optogenetic inhibition. **(c)** Distributions of stably-recorded V1 neurons which were significantly inhibited, activated following light pulses, or unaffected by light delivery (*n* = 58 neurons from 5 mice). **(d)** Power spectra for V1 LFPs showed no significant change in rhythmic activity during periods of inhibition (*N.S*., K-S test, *n* = 5 mice). **(e-f)** During optogenetic inhibition, neurons that showed no decrease in firing rate showed no change in orientation preference (*N.S*., nested t-test, *n* = 32 neurons from 5 mice). In contrast, neurons that were inhibited showed a reduced preference for gratings of the same orientation (X°) used for TRAP (** indicates *p* = 0.007 [*t* = 3.65, *DF* = 8, nested t-test, *n* = 26 neurons from 5 mice).

We next asked whether sleep-targeted inhibition of V1 visual engram neurons (i.e., those encoding the fear memory cues) disrupts consolidation of visually-cued fear memory. For these experiments, *cfos::Arch* mice expressing ArchT in X°-activated neurons (and control mice not expressing ArchT) underwent visually-cued fear conditioning in context A at ZT0, using either X° or Y° as a cue for foot shock (**Figure 7a**). They were then returned to their home cages for *ad lib* sleep. For the first 6 h following conditioning (a window of time where SD disrupts consolidation; **Figure 1**), TRAPed neurons in V1 were optogenetically inhibited (using the parameters described for **Figure 6** above) during bouts of NREM and REM sleep (**Extended Data Figure S3**). This pattern of inhibition did not significantly alter either sleep architecture or V1 EEG power spectra (which were similar between inhibited and control mice; **Extended Data Figure S4**).

**Figure 7.**
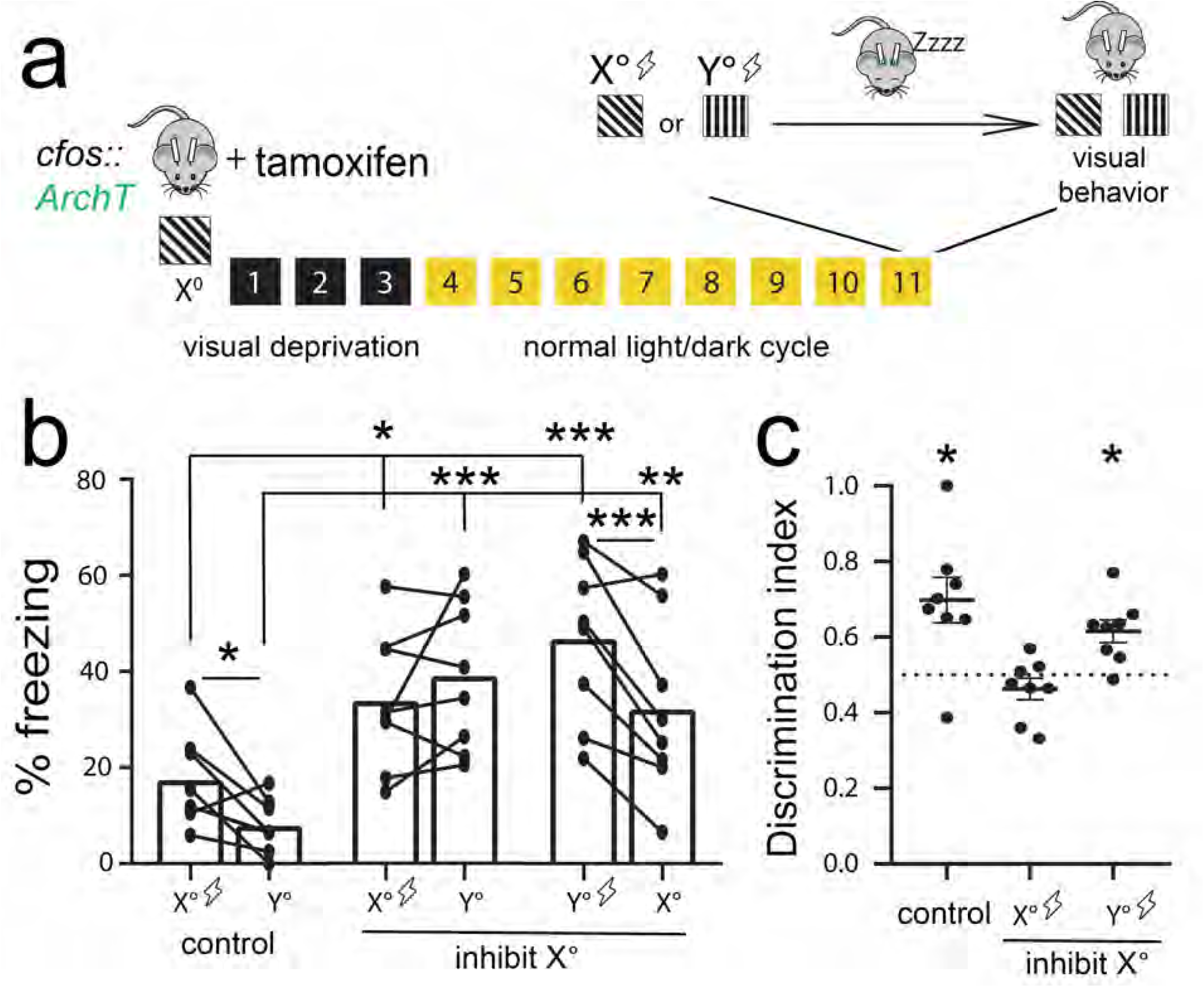
Sleep-specific inhibition of a V1 engram disrupts visually-cued fear memory consolidation. **(a)** *cfos::ArchT* mice implanted with bilateral V1 optical fibers and EEG/EMG electrodes were presented with X° for TRAP. 11 days later, mice were conditioned using either the same orientation (X°) or an alternate orientation (Y°) as the shock cue. Post-conditioning, the mice slept, with sleep-specific inhibition during the first 6h. **(b)** No-inhibition (non-opsin-expressing) controls and mice cued to Y° with subsequent optogenetic inhibition showed higher freezing responses to the shock cue vs. the neutral cue (two-way RM ANOVA: main effect of optogenetic manipulation condition, *F* = 9.9, *p* < 0.001, main effect of orientation, *F* = 9.0, *p* = 0.007, optogenetic condition x orientation interaction, *F* = 7.9, *p* = 0.003, no-inhibition control - *p* = 0.02, Y°-cued inhibition - *p* < 0.001, Holm-Sidak *post hoc* test). In contrast, mice cued to X° with subsequent optogenetic inhibition did not differ in freezing responses to the shock cue vs. the neutral cue (*N.S*., Holm-Sidak *post hoc* test). Mice cued to either X° or Y° with subsequent inhibition showed higher freezing responses to both cues relative to no-inhibition controls, indicative of generalization. (c) Controls and mice cued to Y° show significant discrimination, while mice cued to X° did not (* indicates *p* = 0.016 for both no-inhibition control and Y°-cued with inhibition; Wilcoxon signed rank test).

At ZT12, mice were presented with X° and Y° oriented gratings (shock and neutral cues) in a dissimilar context B. In mice presented with X° as a cue for foot shock, sleep-targeted optogenetic inhibition of TRAPed V1 neurons prevented fear discrimination between X° and Y° cues during testing. These mice showed high levels of generalized fear (i.e., high levels of freezing in response to presentation both gratings) - indicating disrupted fear memory consolidation. In contrast, both control mice (not expressing ArchT) and mice presented with Y° as the shock cue showed cued fear memory consolidation and discriminated between shock and neutral cues at ZT12 (**Figure 7b-c**). Together these data suggest that selective reactivation of V1 visual engram neurons during post-learning sleep provides an essential substrate for consolidation of an associative visually-cued memory.

## Discussion

Our present data demonstrate that orientation-selective V1 neurons involved in encoding a specific visually-cued fear memory (visual engram neurons) play an ongoing role in memory consolidation during subsequent sleep. After selective activation of these neurons during visually-cued fear conditioning, these neurons continue to be active during sleep in the subsequent hours (**Figure 4**) - a time window during which sleep plays a role in promoting consolidation (**Figure 1**). Activity in these neurons is sufficient to drive a percept which can substitute for the visual fear cue in mice during wake (**Figure 3**). It remains unclear how selective sleep-associated reactivation of these neurons affects the surrounding visual cortex (or interacts with circuitry engaged selectively by aversive conditioning). However, periodic optogenetic activation of these orientation-selective neurons is sufficient to drive shifts in orientation preference in surrounding neurons that show excitatory postsynaptic responses to their input (**Figure 5**). This leads to an increase in the representation of the visual engram neurons’ preferred orientation in the surrounding V1 network. While the functions of such an increase in representation are currently unknown, this increase in representation for a specific orientation is seen in the visual cortex in mice^20,27,28,35,36^, human subjects^37,38^, and nonhuman primates^39,40^ as a result of orientation-specific experience and task training. Thus changes in representation in sensory cortex appear to be either a correlate, or a cause, of changes in orientation discrimination ability with experience.

We show conversely, that optogenetic inhibition of orientation-selective neurons acutely reduces the representation for the visual engram neurons’ preferred orientation in the surrounding V1 network (**Figure 6**). Finally, we demonstrate that optogenetic inhibition of these visual engram neurons during post-conditioning sleep dramatically disrupts consolidation of fear memories for specific visual cues (**Figure 7**). Mice with sleep-targeted inhibition of cue-activated neurons show high levels of general freezing behavior at testing, but no discrimination between cues of different orientations. Thus their specific memory deficit seems to be due to an inability to link fear memory to a specific orientation cue during consolidation, rather than a disruption of fear memory *per se*.

This work links together two bodies of literature regarding the neural substrates of memory. One recent area of investigation has focused on the role of engram neurons which are activated by learning experiences, and whose activation is necessary and sufficient for memory recall^24,25,41^. However, the role these neurons play in the consolidation of memories following learning has been a matter of speculation. Here we show that the neurons engaged during learning play a necessary and instructive role during subsequent sleep. The second body of literature has focused on replay of learning-associated activity patterns in specific neuronal ensembles as a mechanism for sleep-dependent facilitation of memory storage. While the phenomenon of replay during sleep has been widely reported^2,7,21,22,42^, a causal role for sleep-dependent replay in memory consolidation has been difficult to prove. At least two technical obstacles have slowed progress toward understanding the role of replay in sleep-dependent consolidation. First, many tasks used in rodents to study phenomena (e.g. maze running) that require several days of training prior to obtaining recordings of sequential firing patterns - a timescale incompatible with memory consolidation occurring across a single sleep period. Second, many prior studies aimed at addressing the question of replay’s necessity for consolidation have relied on disrupting circuit-level activity across windows of sleep^18,19^, sometimes over several days^16,17^. Here we have taken advantage of recently-developed genetic tools to label cue-activated neurons^33^ and a new single trial paradigm for studying sleep-dependent consolidation of memory for a specific sensory cue (**Figure 1**). These have allowed us to demonstrate that sleep-associated reactivation of cue-activated visual engram neurons plays a critical, instructive role in consolidating an associative memory linked to that cue.

A limitation of the present study is that inhibition of visual engram neurons in V1 occurred throughout all stages of sleep (i.e., both REM and NREM). Our prior work on experience-dependent plasticity in V1 has demonstrated that thalamocortical oscillations coordinating activity between the lateral geniculate nucleus (LGN) and V1 during NREM sleep are essential for orientation preference shifts in V1^20^. The pattern of optogenetic stimulation used on visual cue-activated neurons in this study (i.e., regular periodic activation at 1 Hz) is in some ways similar to what occurs in V1 during these NREM oscillations. Critically, this pattern of activation is sufficient to drive large V1 orientation preference shifts (**Figure 5**). However, a role for REM activity in cortical plasticity cannot be ruled out. REM plays a critical role in developmentally-regulated experience-dependent plasticity in V1^43^. In many species, REM is characterized by selective activation of LGN-V1 circuitry during pontine-geniculate-occipital (PGO) waves, which promote synaptic plasticity in various brain structures^14^. Future work will be aimed at both characterizing patterns of activity in orientation-selective populations during REM vs. NREM, and in targeting inhibition of this population to specific states.

The present findings may ultimately inform our understanding of how sensory cortical areas interact with structures such as the hippocampus and amygdala during sleep, and how these interactions inform consolidation of specific memories. Together our data indicate that primary sensory structures engaged in fear memory encoding communicate with structures conveying emotional valence information during post-learning sleep to promote long-lasting fear association with a specific cue. Whether this interregional communication is unique to one or more sleep states is a critical unanswered question. Answering this question may have important implications not only for understanding sleep’s mechanistic role in memory consolidation, but also its mechanistic role in regulation of mood and affect. It will also have specific implications for treating disorders where fear is dysregulated or misattributed, including anxiety and panic disorders, acute stress disorder, and PTSD.

## Materials and Methods

### Animal handling and husbandry

All animal procedures were approved by the University of Michigan Institutional Animal Care and Use Committee. With the exception of constant dark following tamoxifen administration, mice were kept on a 12 h:12 h light:dark (LD) cycle, and were given food and water *ad lib* throughout the entirety of the study. Following surgical procedures, and during habituation prior to cued conditioning, mice were individually housed in standard caging with beneficial environmental enrichment (nesting material, manipulanda, and/or novel foods).

### Visually-cued fear conditioning

For 3 days prior to conditioning, mice were habituated to 5 min/day of gentle handling. Following the habituation period, at ZT0, mice underwent visually-cued fear conditioning in a novel arena (context A). They were allowed 2 minutes to acclimate to the arena. They then experienced 3 pairings of a 30-s visual stimulus (presented simultaneously on 4 LED monitors surrounding the arena) co-terminating with a 2-s 0.75 mA foot shock. These pairings were divided with a 60-s intertrial interval. Each visual stimulus consisted of a 1 Hz phase-reversing oriented grating (X°) with a spatial frequency of 0.05 cycles/degree and contrast of 100%.

Following conditioning, C57BL/6J mice (Jackson) used for experiments outlined in **Figure 1** were returned to their home cage and were either allowed 12 h *ad lib* sleep, or were sleep deprived (SD) using gentle handling (i.e., cage tapping, nest disturbance, and light touch with a cotton-tipped applicator to cause arousal from sleep) for 6 h, after which they were allowed 6 h *ad lib* sleep. All transgenic mice (see below) with data shown in **Figures 2**, **3**, and **7** were allowed *ad lib* sleep in their home cage following conditioning.

At ZT12 (i.e., 12 h following conditioning) mice were placed in a dissimilar novel context B for cued fear memory testing. Context B differed from context A (used during conditioning), with the two arenas having a unique odor, shape, size, floor texture, and lighting condition. During testing, mice were exposed to two distinct oriented grating stimuli (X° and Y°) to assess cue discrimination. At the start of each test, mice were allowed 3 min to acclimate to the arena, after which an oriented grating either the same as the shock cue (X°) or distinct (Y°) was presented for 3 min followed by 1 min of post stimulus arena exploration. A minimum of thirty minutes were left between the presentations of the tests.

Freezing responses were quantified for each grating stimulus using previously-established criteria^44^. For each test, two scorers blinded to behavioral condition quantified periods of immobility during presentation of grating stimuli that included fear features such as hyperventilation and rigid posture. Freezing during presentation of the two gratings was compared to calculate a discrimination index: (percent freezing during shock [X°] simulus)/(percent freezing during shock [X°] stimulus + percent freezing during neutral [Y°] stimulus).

To test for time-of-day effects on visually-cued fear memory recall (**Supplemental Figure 2**), additional cohorts of mice were trained at ZT0 as described above, and tested at 12, 24, or 36 h later.

### Genetic tagging of orientation-selective V1 neurons

Prior to all procedures for targeted recombination in visual engram neurons, mice were habituated for 3 days to gentle handling procedures. After habituation, at ZT0, the mice were placed in this square arena surrounded by 4 LED monitors. Each monitor presented a single-orientation (X°) phase-reversing grating stimulus (1 Hz, 0.05 cycles/degree, 100% contrast) for 30 min (or, for negative controls shown in **Figure 2**, a dark screen). Immediately after stimulus or dark screen presentation, mice received an i.p. injection of tamoxifen (100mg/kg in 95% corn oil/ 5% ethanol), and were placed in complete darkness for the next 3 d to prevent further visually-driven recombination in V1. Following 3 d of constant dark housing, mice were returned to a normal 12 h:12 h LD cycle for 7 d prior to further experiments. cfos-CRE^ER^ mice (Guenthner et al 2013; B6.129(Cg)-Fos^*tm1.1(cre/ERT2)Luo*^/J; Jackson) crossed to either B6.Cg-*Gt(ROSA)26Sor^tm9(CAG-tdTomato)Hze^*/J, B6.Cg-*Gt(ROSA)26Sor^tm32(CAG-COP4^*^H134R/EYFP)Hze^*/J, or B6.Cg-*Gt(ROSA)26Sor^tm40.1(CAG-aop3/EGFP)Hze^*/J (Jackson) mice to induce CRE recombinase-mediated expression of tdTomato, ChR2, or ArchT.

### Histology and immunohistochemistry

At the conclusion of each experiment, mice were deeply anesthetized with pentobarbital, and transcardially perfused with saline and 4% paraformaldehyde. Brains were dissected, post-fixed, cryoprotected in 30% sucrose, and cryosectioned at 50 μm. Transgene expression in V1 was verified for all experiments using CRE-dependent transgenic lines prior to subsequent data analysis. For electrophysiological recordings in V1, electrode placement was verified prior to data analysis. Immunohistochemistry for cFos was carried out using rabbit-anti-cfos 1:1000 (Abcam; ab190289) and secondary donkey-anti-rabbit conjugated to Alexa Fluor 405 (1:200; Abcam; ab175651); coronal sections containing V1 were mounted using Fluoromount-G (Southern Biotech). Co-labeling of tdTomato and anti-cFos was quantified using Image J software in 6 sections containing V1 from each mouse by a scorer blinded to animal condition. Average co-labeling values for each mouse are reported in **Figures 2** and **4**.

### V1 visual response recordings, optogenetic manipulations, and data analysis

For anesthetized recordings of V1 neurons’ visual responses and firing, mice were anesthetized using a combination of isoflurane (0.5-0.8%) and 1 mg/kg chlorprothixene (Sigma). Data was acquired using a 32-channel Plexon Omniplex recording system, using previously-described methods^20,22^. A 2-shank, linear silicon probe (250 μm spacing between shanks) with 25 μm inter-electrode spacing (16 electrodes/shank; Cambridge Neurotech) was slowly advanced into V1 until stable recordings (with consistent spike waveforms continuously present for at least 30 min prior to baseline recording) were obtained. Orientation tuning curves for recorded neurons were generated by presenting a series of 8 full-field phase-reversing oriented gratings (0, 22.5, 45, 67.5, 90, 112.5, 135, or 157.5 degrees from horizontal, 1 Hz, 0.05 cycles/degree, 100% contrast, 10 s duration) and a blank screen (to evaluate spontaneous activity) presented repeatedly (4-8 times each) in an interleaved manner.

For recordings during rhythmic optogenetic activation of X°-activated V1 neurons in ChR2-expressing mice (**Figure 5**) tuning curves were generated: 1) at baseline, 2) after a 20-30 min period without optogenetic manipulation, and 3) after a 20-30 min period of 1 Hz optogenetic stimulation. Optogenetic stimulation consisted of blue light pulses (10 ms, 473 nm, 10 mW power) delivered at 1 Hz. Only neurons stably recorded throughout all phases of the experiment (shown in **Figure 5A**) were included in firing and visual response analysis.

To assess effects of optogenetic inhibition of X°-activated V1 neurons in ArchT-expressing mice (**Figure 6**) recordings consisted of a 30-min spontaneous activity recording with no manipulation, a 30-min recording with periodic inhibition (532 nm green light, 15 mW, delivered in cycles of 5 s on, followed by a 500 ms offramp and a 1-s off period). Following these recordings, two orientation tuning curves were generated for all recorded neurons: 1) a baseline without inhibition, and 2) with inhibition of X°-activated V1 neurons occuring during 10-s presentations of oriented grating stimuli. Only neurons stably recorded throughout all phases of the experiment (shown in **Figure 6A**) were included in firing and visual response analysis.

For all recordings, stable single units were isolated using PCA-based analysis and MANOVA-based cluster separation, implemented using Offline Sorter software (Plexon) and previously-described methods^20,22^. Units that could not be reliably discriminated, or had refractory period violations in their spiking patterns, were eliminated from subsequent analyses. Changes in orientation tuning were assessed relative to the orientation of the TRAPed ensemble (X°), based on neurons mean firing rate responses to gratings of different orientations. For each tuning curve, an orientation preference index (OPI) was calculated for X° and the orthogonal stimulus orientation (X°/X+90°), as described previously^20,27,28^. % changes in OPI (across optogenetic stimulation or control conditions) were calculated as [(OPI^pre^-OPI^post^)/OPI^pre^] *100. Firing responses of neurons during rhythmic optogenetic stimulation in ChR2-expressing mice was assessed from Z-scored perievent rasters centered on blue light onset; significance of time-locked excitation or inhibition was calculated based on positive or negative Z-score deviations beyond the 99% confidence interval (Neuralynx; Plexon). Changes in firing during optogenetic inhibition in ArchT-expressing mice were calculated for each neuron within the inhibition recording period, by comparing mean firing rate during the last 1.5 s of each green light delivery period with mean firing rate during the subsequent 500 ms offramps and 1-s off period.

Power spectral density for local field potentials was detrended using NeuroExplorer software (Plexon) with a single taper Hann Windowing Function with 50% window overlap. These were averaged across all active electrodes on each silicon probe shank. Distributions of power (between 0 and 20 Hz) were compared statistically using KS tests.

### Surgical procedures

For V1 optical fiber implantation, mice were anesthetized using 1-2% isoflurane. Optical fibers (0.5 NA, 300 um core, ThorLabs) were positioned bilaterally at the surface of V1 at a 80 degree angle relative to the cortical surface (2.9 mm posterior, 2.7 mm lateral). Implants were secured to the skull with an anchor screw positioned anterior to bregma, using Loctite adhesive. For EEG/EMG recordings to differentiate sleep states, in addition to bilateral V1 optical fibers, mice received an EEG screw over V1 (2.9 mm posterior, 2.3 mm lateral), a reference screw over the cerebellum, and an additional EMG electrode in nuchal muscle. Mice were allowed 10 days of postoperative recovery before procedures to induce transgene expression in V1.

### Optogenetic manipulations in behaving animals

Two cohorts of implanted mice, expressing ChR2 in the TRAPed ensemble, were used to test perception of optogenetic activation of this cell population. Prior to behavioral training and testing, these mice were habituated to handling and tethering (for light delivery to V1) procedures for 3 days. The first cohort underwent cued fear conditioning as described above in context A at ZT0, with 30-s blocks of rhythmic light delivery to V1 (1 Hz, 10 mW, 10 ms pulses) serving as a proxy shock cue (i.e., substituting for visual oriented grating presentation). Following 3 optogenetic stimulation-shock pairings, these mice were returned to their home cages and allowed *ad lib* sleep until ZT12. At ZT 12, they were placed in context B and freezing responses were assessed for visual presentation of both the same orientation as the TRAPed ensemble (X°) and an alternate orientation (Y°), as described above. A second cohort of mice underwent visually-cued fear conditioning to the same angle as the TRAPed ensemble (X°) in context A at ZT0. After conditioning, they were returned to their home cage for *ad lib* sleep. At ZT12, they were tested in context B, where freezing behavior was assessed before, during, and after a period of 1 Hz light delivery to V1 (3 min before, 3 min during, and 1 min after).

To assess effects of sleep-targeted inhibition of visual engram neurons, 10 days after EEG/EMG and optical fiber implantation, mice underwent procedures to induce expression of ArchT in the TRAPed orientation-specific ensemble. Following 3 days of habituation to handling and tethering (for light delivery to V1 and EEG/EMG recording), these mice underwent 12 h sleep/wake baseline recordings, starting at ZT0. The next day, mice underwent visually-cued fear conditioning at ZT0, using either the same orientation as the TRAPed ensemble (X°) or an alternate orientation (Y°) as a cue for foot shock. They were then returned to their home cage for *ad lib* sleep. For the first 6 h post-conditioning, a subset of mice expressing ArchT underwent periodic optogenetic inhibition targeted to both NREM and REM sleep. The state targeting was based on EEG signals, EMG signals, and the animal’s behavior. A control group of mice which were not expressing ArchT underwent the same light delivery and recording procedures. At ZT12, all mice were placed in context B to assess freezing responses to both X° and Y° oriented gratings, as described above.

EEG and EMG signals were used offline to classify each 10-s interval of baseline and post-conditioning recording periods as either wake, NREM, or REM sleep, using custom MATLAB software^20,22^. Additionally microarousals (periods of non-oscillatory activity between periods of NREM) as small as 5 s were identified as wake. Mean power spectral density was calculated separately within REM, NREM, and wake for each phase of recording, and within and outside of periods of light delivery to V1, as described previously^20^. The power spectra were calculated as percent of the total spectral power.

### Statistical methods

All statistical analyses were done using GraphPad Prism. Prior to making comparisons across values, the normality of distributions was tested using the D’Agostino-Pearson omnibus k2 test. Nonparametric tests were used when data distributions were non-normal or when *n* values were too low to test normality. If the data involved multiple data measurements from one animal (e.g. multiple images taken from the same animal for immunohistochemistry), nested statistics were used. All statistical tests were two-tailed. For each specific data set the statistical tests used are listed in the **Results** section. *p*-values are represented as * *p* < 0.05, ** *p* < 0.01, *** *p* < 0.001, \*\*\*\**p* < 0.0001

## Data availability

All relevant data and analysis tools are available upon reasonable request from the authors.

## Code availability

Any MATLAB codes used in analysis are available from the authors upon reasonable request.

## Acknowledgements

The authors are grateful to members of the Aton lab, and to Drs. Natalie Tronson, Monica Dus, Richard Hume, and Dawen Cai for helpful feedback on this manuscript. This work was supported by research grants from the NIH (R01 NS104776) and the Human Frontiers Science Program (N023241-00_RG105) to SJA, a NSF Graduate Research Fellowship to BCC, and a Rackham Graduate Fellowship to BCC.

## Extended Data Figure Legends

**Figure S1.**
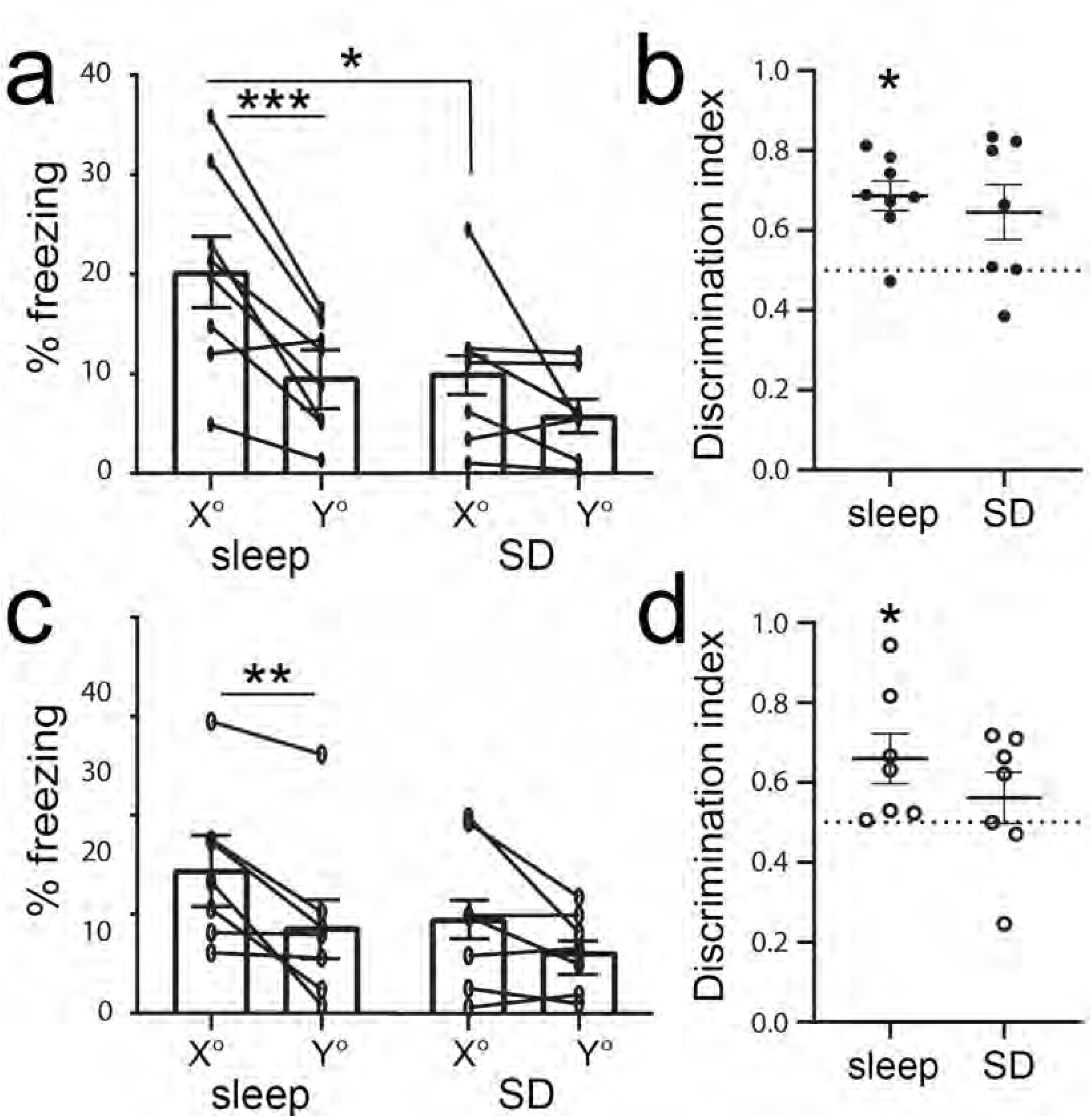
Both female and male mice show deficits in visually-cued fear memory following post-conditioning sleep deprivation. **(a)** Male mice allowed *ad lib* sleep following conditioning froze significantly more to the shock cue (X°) than mice who were sleep deprived (* indicates *p* = 0.01, Holm-Sidak *post hoc* test). Freely sleeping male mice froze significantly more in response the shock cue than a neutral cue (*** indicates *p* = 0.001, Holm-Sidak *post hoc* test). **(b)** Sleeping, but not sleep deprived, male mice showed discrimination for shock vs. neutral cues above chance (* indicates *p* = 0.02, Wilcoxon signed rank test). **(c)** Female mice who were allowed *ad lib* sleep also showed higher freezing responses to the shock cue than the neutral cue (** indicates p = 0.007, Holm-Sidak *post hoc* test). **(d)** Female mice allowed *ad lib* sleep showed discrimination in their responses to shock vs. neutral cues, while sleep deprived female mice did not (* indicates p = 0.02, Wilcoxon signed rank test vs chance).

**Figure S2.**
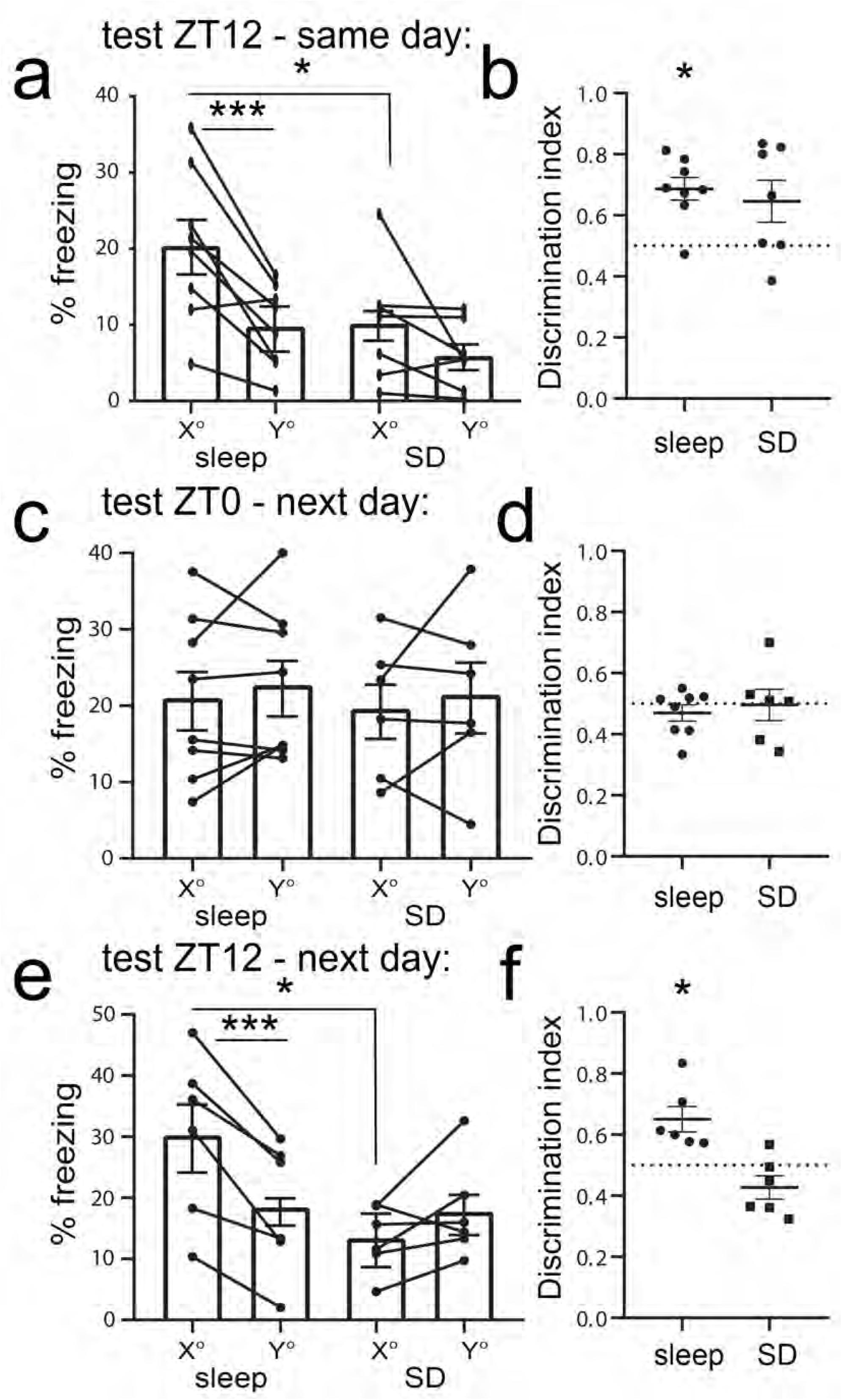
Discrimination of fear cues at testing follows a diurnal pattern. **(a)** ZT12 cued fear test performance, carried out on the day of conditioning. Sleeping mice showed higher freezing in response to the shock cue than SD mice (* indicates *p* = 0.01, Holm-Sidak *post hoc* test). Sleeping mice froze more in response to presentation of the shock cue than the neutral cue (*** indicates *p* = 0.001, Holm-Sidak *post hoc* test). **(b)** Sleeping mice, but not SD mice, showed freezing response discrimination between shock and neutral cues (* indicates *p* = 0.02, Wilcoxon signed rank test vs chance) **(c)** ZT0 cued fear test performance, carried out 24 h following conditioning. There were no differences within groups or across groups (*N.S*., Holm-Sidak *post hoc* test). **(d)** Neither group discriminated between shock and neutral cues beyond chance (*N.S*., Wilcoxon signed rank test vs chance). **(e)** ZT12 cued fear test performance, carried out on the day following conditioning. Freely sleeping mice showed greater freezing in response to the shock cue than SD mice (* indicates *p* = 0.01, Holm-Sidak *post hoc* test). Sleeping mice froze more in response to presentation of the shock cue vs. the neutral cue (*** indicates *p* < 0.001, Holm-Sidak *post hoc* test). **(f)** Mice allowed *ad lib* sleep (but not SD mice) showed discrimination of freezing responses to shock vs. neutral cues above chance (* indicates *p* = 0.03, Wilcoxon signed rank test vs. chance).

**Figure S3.**
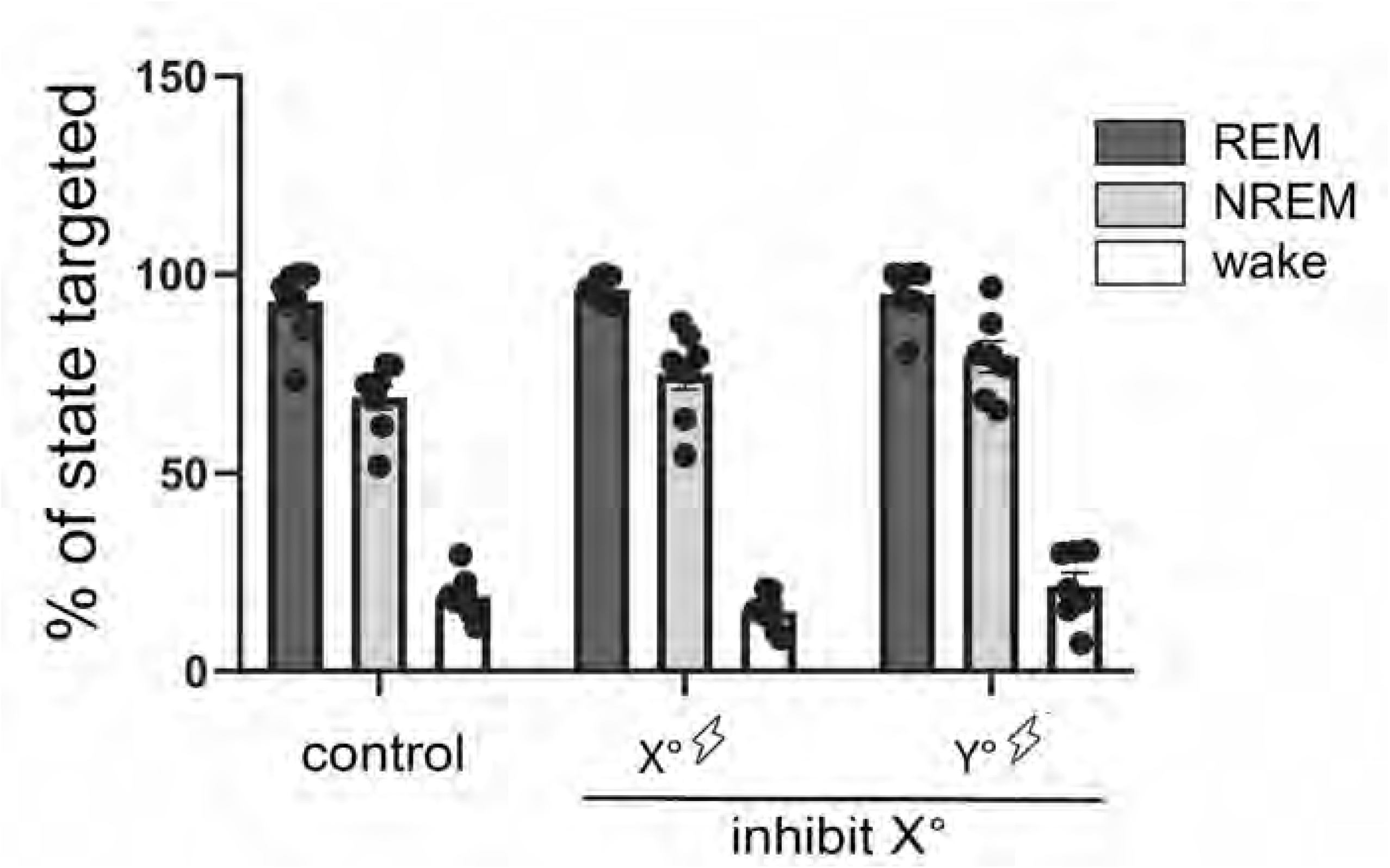
State-specific targeting of optogenetic inhibition. There were no significant differences in state coverage between different experimental groups (*N.S*., two-way RM ANOVA for *n* = 8 no-opsin [no-inhibition] control mice, *n* = 8 mice cued to X° with subsequent inhibition, *n* = 7 mice cued to Y° with subsequent inhibition). In each group, light was delivered to V1 throughout most of REM sleep (93 ± 3%, 96 ± 1%, and 96 ± 3% of total REM, respectively) and NREM sleep (69 ± 3%, 75 ± 4%, and 79 ± 3% of total NREM, respectively) were covered. In each group there was a small amount of light delivery during wake, primarily during microarousals (19 ± 2%, 15 ± 2%, and 22 ± 3% of total wake, respectively).

**Figure S4.**
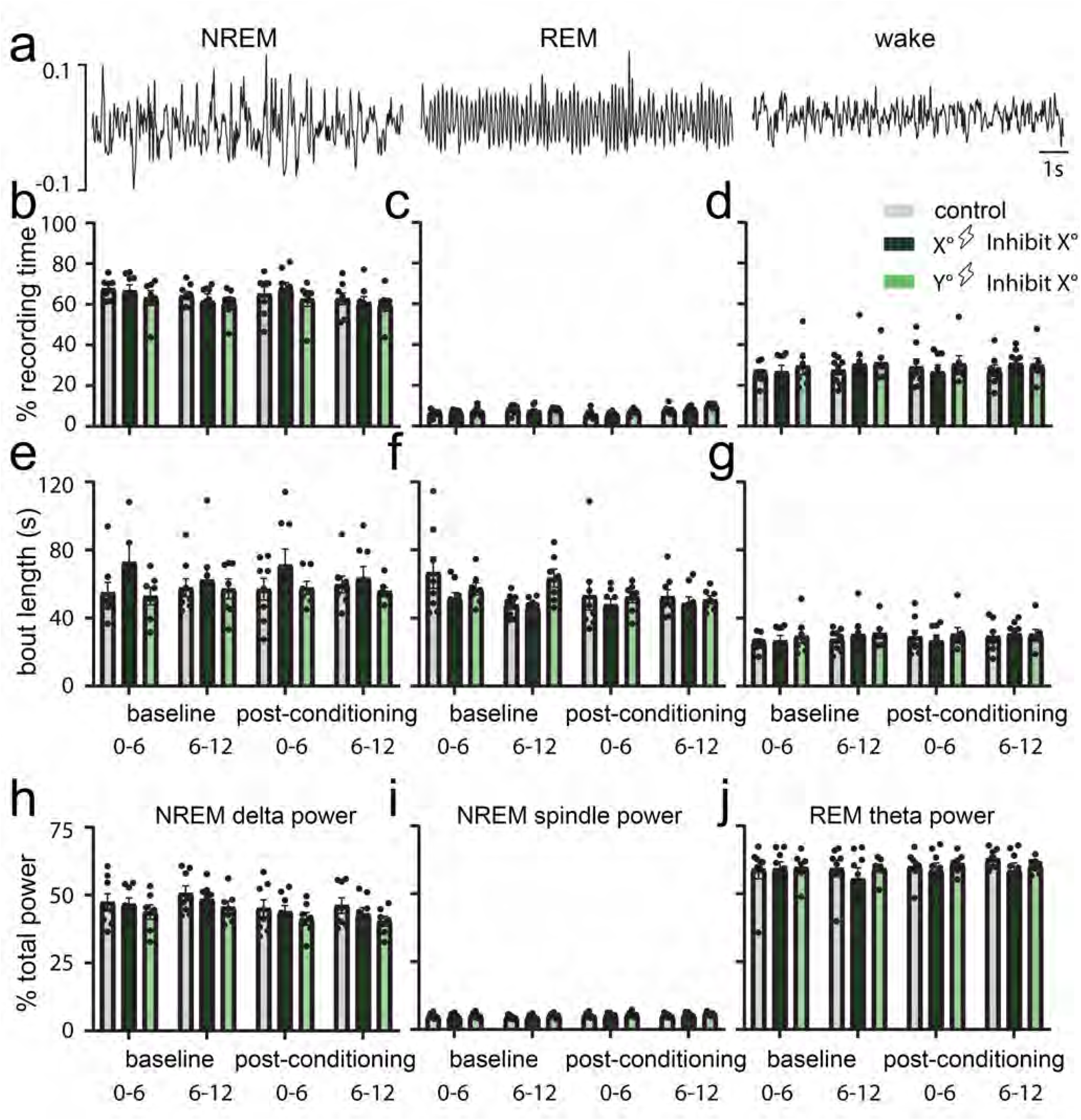
Sleep architecture and power during baseline and optogenetic inhibition. (a) Representative traces of EEG classified as NREM sleep, REM sleep, and wake. (b-d) Percent of recording time spent in each state across recording periods and experimental groups. There were no significant differences in sleep time between groups (*N.S*., two-way RM ANOVA). (e-g) Average bout length for each state across recording times and across experimental groups. There were no significant differences between groups (*N.S*., two-way RM ANOVA). (h-j) Average power within NREM delta (0.5-4 Hz), NREM spindle (12-15 Hz), and REM theta (4-12 Hz) frequency bands across recording periods and experimental groups. There were no significant differences between groups (*N.S*., two-way RM ANOVA).

